# Isolation and biophysical characterization of GSU0105, a triheme c-type cytochrome from *Geobacter sulfurreducens*

**DOI:** 10.1101/2020.11.03.367284

**Authors:** Tyler J. Brittain, Matthew C. O’Malley, Coleman M. Swaim, Reilly A. Fink, Oleksandr Kokhan

**Affiliations:** Department of Chemistry and Biochemistry, James Madison University, Harrisonburg, VA 22807 USA

**Keywords:** *Geobacter*, Fe(III) respiration, Multiheme cytochromes, Electron transfer

## Abstract

C-type cytochromes play an important role in respiration of dissimilatory metal-reducing bacteria. They form extended conduits for charge transfer between the cellular metabolism and external electron acceptors such as particles of iron oxide, metal ions, and humic substances. Out of more than a hundred c-type cytochromes in *Geobacter sulfurreducens*, only a small fraction has been previously characterized. Here we present our results on expression and biophysical characterization of GSU0105, a novel 3-heme cytochrome, important for Fe(III) respiration in *G. sulfurreducens*. We successfully cloned the gene and achieved ~3 mg/L of culture GSU0105 expression in *E.coli*. Despite a similar size (71 amino acids) and the same number of c-type hemes to the members of the cytochrome (cyt) c_7_ family, multiple sequence alignment suggests that GSU0105 does not belong to the cyt c_7_ family. UV-Vis spectroscopy revealed typical c-type cytochrome spectral features, including a weak iron-sulfur charge transfer band suggesting that at least one heme is ligated with a methionine residue. Far UV circular dichroism studies demonstrate approximately 35% content of α-helices and β-sheets, each, as well as thermal aggregation occurring above 60 °C. A combination of SAXS and analytical size exclusion chromatography data shows that GSU0105 is monomeric in solution. Finally, affinity pull-down assays demonstrate high binding affinity to PpcD and weaker binding to the other members of the cyt c_7_ family.

## 1. Introduction

*Geobacter sulfurreducens* can utilize Fe(III) and Mn(IV) oxide particles[1–3], U(VI)[4], and humic substances[5] for its respiration. This makes it an attractive target for bioremediation applications and for electricity generation with bacterial fuel cells[6–12]. *G. sulfurreducens’* genome encodes 111 c-type cytochromes[13]. The evolutionary need for such a large number of cytochromes is not clear and only a small subset has been biochemically and biophysically characterized. The vast majority of the previous work was focused on five members of the triheme cyt c_7_ family, PpcA-E[14–22]. More limited data is available for OmcF[23–26], OmcS[27, 28], OmcZ[29, 30], and a few other cytochromes[31–34].

One of the largest up-to-date studies of cytochrome expression in *G. sulfurreducens* under various growth conditions utilized 2-D PAGE with LC-MS and identified 91 out of 111 predicted cytochromes[35]. One of the most interesting results of the study was the identification of three cytochromes (GSU0105, GSU0701, GSU2515) present in cultures grown under Fe(III) reducing conditions, but not expressed in cultures grown on fumarate. The largest of these three proteins, GSU0701 (also known as OmcJ) is expected to have 6 heme binding sites and to be localized on the outer membrane as a member of the Outer Membrane Cytochrome (Omc) family. Based on the sequence analysis, GSU2515 has one c-type heme. It is expected to be localized either in the periplasm or on the outer membrane[36]. Finally, GSU0105 is expected to have three c-type hemes and is likely to be located in the periplasm[37] to serve as an electron carrier linking cytochromes on the outer membrane with intracellular respiration when cells are grown under Fe(III) reducing conditions.

Despite GSU0105’s central role in Fe(III) respiration, this protein has not been previously characterized. Some preliminary data was reported in a M.S. thesis by T.M. Fernandes[37], but a low yield of the recombinant protein in *E. coli* (~0.16 mg/L) seemed to preclude the author and his co-workers from performing a comprehensive study. Our report demonstrates a different construct for recombinant GSU0105 expression with much higher yields, which allowed us to perform the comprehensive biophysical characterization described in this report.

## 2. Materials and methods

### 2.1. Reagents

Genomic DNA from *G. sulfurreducens* was a generous gift from Prof. D. R. Lovley (University of Massachusetts-Amherst). PCR primers for gene cloning were synthesized by Invitrogen (Carlsbad, CA). Q5 PCR master mix, T4 ligase, T4 kinase, and DpnI were purchased from New England Biolabs (Ipswich, MA). Ampicillin, carbenicillin, chloramphenicol, and IPTG were purchased from Gold Bio (St. Louis, MO). Sucrose, cholate, agar, and agarose were from Bio Basic (Amherst, NY). Macro S cation exchange resin was purchased from BioRad (Hercules, CA). Proteins for size-exclusion column calibration were purchased from Sigma-Aldrich (St. Louis, MO). 5-aminolevulinic acid was purchased from Carbosynth (San Diego, CA). All other reagents, unless otherwise specified, were purchased from Fisher Scientific (Waltham, MA).

### 2.2. Cloning of GSU0105

For an unrelated project in our lab we previously modified a pVA203 vector[31] by inserting a Factor Xa cleavable His-tag sequence (IDGRHHHHHH) at the C-terminus of the PpcA gene. The sequence of this new plasmid (pVA203XaHT) was verified at the University of Chicago DNA Sequencing and Genotyping Facility. We predicted the *G. sulfurreducens* periplasmic signaling sequence in GSU0105 (GenBank: AAR33440.2) using SignalP 5.0[38]. We used restriction-free cloning to replace the PpcA gene sequence in the pVA203XaHT vector with the GSU0105 gene. During the cloning step we deleted the native GSU0105 signaling sequence from the GSU0105 gene and cloned the product between the OmpA signaling sequence and the cleavable His-tag sequences present in pVA203XaHT. PCR primers (5’-<u>TTTCGCTACCGTTGCGGCCGCAGCCTTCGAATGCAACGTC</u>-3’ and 5’-<u>GGTGGTGGTGACGGCCATCAATTTCTCCAGCCTGCGGTTT</u>-3’) for restriction-free cloning of GSU0105 from genomic *G. sulfurreducens* DNA were designed using RF-Cloning[39]. Both PCR steps followed the recommended temperatures and conditions using Q5 Master Mix (New England Biolabs). The PCR products from the first step were purified using 1% agarose gel. The band corresponding to the GSU0105 gene with overhangs for the second step of PCR was excised from the gel and purified using an EZ-10 Gel Extraction miniprep kit (BioBasic). The double-stranded DNA segments produced in the previous step were used as primers for the second PCR reaction with the pVA203XaHT vector as the template to insert the GSU0105 gene in place of PpcA. The resulting PCR product was purified with an EZ-10 PCR Product miniprep kit (BioBasic) and treated with DpnI to remove the template DNA. Finally, the product was phosphorylated and ligated to create a new vector pGSU0105XaHT. This vector was transformed into chemically competent 5-α *E. coli* cells (New England Biolabs), outgrown for 1 hour on SOC medium and plated on LB agar plates supplemented with 100 mg/L of carbenicillin. The plates were incubated overnight at 37°C. Individual colonies were picked and grown on LB medium supplemented with 100 mg/L ampicillin. Plasmid DNA was extracted using an EZ-10 Plasmid DNA miniprep kit (Biobasic) and the cloned gene sequence was verified at the University of Chicago DNA Sequencing and Genotyping Facility.

### 2.3. Protein Expression and purification

GSU0105 was expressed in the BL21 (DE3) *E. coli* strain containing *c*-type cytochrome maturation genes (Ccm) in the pEC86 vector[40]. The cells were grown on 2xYT medium at 28 °C. After the cell culture absorbance reached 1.5 OD at 600nm, the cells were induced with 20 μM IPTG and 1 mM 5-aminolevulinic acid (heme biosynthesis precursor) and grown for additional 14-16 hours at 28 °C. The harvested cell pellets were frozen at −80 °C. The first step of GSU0105 purification was similar to our PpcA isolation procedure[41]. In short, cell pellets were thawed and resuspended in 1:20 (w/v) TSE buffer (20 mM Tris, pH 7.5, 20% (w/v) sucrose, 0.5 mM EDTA, 40mg/L lysozyme) and gently shaken for 30 min at room temperature. Cell suspensions were centrifuged at 8,000 g for 20 min. The supernatant fraction (“periplasm”) was applied to a Macro S (BioRad) cation exchange column equilibrated with 20 mM Tris, pH 7.5, and 20 mM cholate (“Buffer A”) and washed with at least 5 column volumes of Buffer A. GSU0105 was eluted with Buffer A supplemented with 200 mM NaCl. The eluted colored fraction was applied to a Ni-NTA column equilibrated with Buffer A and 200 mM NaCl. The column was washed with at least 5 column volumes of the same buffer. GSU0105 was eluted with Buffer A supplemented with 200 mM NaCl and 300 mM imidazole. GSU0105 was then concentrated and buffer exchange was preformed to Buffer A with Amicon Ultra-15 filter units (3 kDa cut-off). Finally, the concentrated GSU0105 samples were centrifuged at 15,000 *g* for 15 minutes to remove aggregates.

PpcA was isolated as described in [41]. PpcB, PpcC, PpcD and PpcE were expressed and isolated as previously described[16].

### 2.4. Circular dichroism spectroscopy

Circular dichroism (CD) spectroscopy was performed on a temperature-controlled Jasco J-1500 CD spectrometer with 14.8 μM GSU0105 in 1 mm quartz cuvettes in Buffer A. Triplicate spectra were recorded between 190 and 350 nm and averaged to improve the signal-to-noise ratio. The buffer and cuvette background spectra were subtracted and protein secondary structure was predicted using BeStSel[42].

### 2.5. Analytical size-exclusion chromatography

Analytical size exclusion chromatography was performed with 20μl protein samples loaded on a Superdex 200 Increase 10/300 GL column (GE Healthcare) mounted on an Agilent 1260 HPLC. The running buffer contained 20 mM Tris, pH 7.5, 200 mM NaCl, and 20 mM cholate. The separation was performed at 0.75 ml/min flow rate. Eluted proteins were detected with a diode array spectrophotometer recording absorbance values at 210, 280, 408 nm. Calibration of the size exclusion column was performed with 20-50 μl injections of 10-20 mg/ml solutions of PpcA (9.6 kDa), bovine serum albumin (66.6 kDa), bovine blood γ-globulin (150 kDa), equine spleen ferritin (440 kDa), or bovine thyroglobulin (669 kDa).

### 2.6. Small-angle X-ray scattering

Small-angle X-ray scattering (SAXS) was performed at 12-ID-B beamline of the Advanced Photon Source at Argonne National Lab as previously reported[41]. In short, ~100 μM His-tag and Factor Xa cleaved GSU0105 protein samples in 20 mM Tris, pH 7.5 were loaded in a quartz flow cell and gently refreshed with a syringe pump during X-ray (14 keV) exposures to minimize protein damage. 30 images with 1 sec exposure times were recorded with a Pilatus 2M detector. The buffer background signal was collected under the same conditions and subtracted from the data for the protein solutions. The momentum transfer values calibration was performed with silver behenate calibration standards.

### 2.7. Affinity pull-down assays

HisPur Ni-NTA slurry (Thermo Scientific) was diluted 5-fold with 20 mM Tris, pH 7.5 buffer (“Buffer B”) and centrifuged for 2 min at 100 *g* to sediment the resin. The supernatant was carefully removed. The pellet was twice more resuspended 1:10 (v/v) in Buffer B and centrifuged to remove any residual ethanol from the storage solution and to equilibrate the resin. After the third washing step, the resin pellet was resuspended in Buffer B and divided into two equal volumes. One volume was incubated with GSU0105 in Buffer B and the other was incubated with a similar volume of Buffer B without any protein. After 30 min incubation with gentle shaking, both resin pellets were sedimented at 100 *g* and washed 3 times with a 5-fold volume excess of Buffer B. After the final wash step, the resin pellets were resuspended again in Buffer B and pipetted into 1.5 mL microcentrifuge tubes. After centrifugation at 100 *g* each tube contained approximately 200 μL of settled resin volume. At this point, the tubes with GSU0105 bound to the resin (“+GSU0105”) had approximately 1.3 nmol of GSU0105 per tube. “-GSU0105” tubes had the same volume of the resin equilibrated in Buffer B without any proteins bound.

This step was followed by additions of 1 mL of 10 μM solutions of Ppc cytochromes in Buffer B to the previously prepared tubes with GSU0105 bound and unbound resin. After 30 min incubation with gentle shaking, the Ni-NTA resin samples were sedimented again by centrifugation for 2 min at 100 *g*. The pellets were washed 3 times with 1 ml of Buffer B and the same centrifugation parameters. After the last wash step, 1 mL of Buffer B with 500 mM imidazole was added. After 10 min incubation, the tubes were centrifuged for 2 min at 1,000 *g* and UV-Vis spectra of supernatants were recorded in 1 cm quartz cuvettes using an Agilent 8453 diode array spectrophotometer to quantify the amounts of eluted cytochromes from their peak absorbances at 408 nm. For each tested cyt c_7_ family member, 3-4 independent experiments were performed.

### 2.8. Other characterization techniques

UV-Vis measurements were performed on an Agilent 8453 diode array spectrophotometer with 1 cm quartz cuvettes. All protein dilutions were made with 20 mM Tris, pH 7.5, 20 mM cholate, unless stated otherwise. All proteins were isolated with fully oxidized hemes, negating the need for the addition of any oxidizing agents. The reduced spectrum was obtained by adding small amounts of sodium dithionite to the cuvette until the 552 nm peak ceased to grow.

GSU0105 heme concentration was quantified with a pyridine hemochromagen assay according to the procedure from Barr and Guo[43] and using the recommended extinction coefficient ɛ = 23.97 mM-1*cm^−1^ for reduced-*minus-*oxidized absorbances at 550 and 535 nm[43, 44]. The GSU0105 extinction coefficient was calculated assuming the presence of 3 hemes per mature cytochrome.

Non-reducing SDS-PAGE analysis was performed with 10% TruPAGE precast gels (Sigma-Aldrich) following the manufacturer protocol with MOPS buffer. Low-range Rainbow molecular weight markers (GE Healthcare) were used to estimate the masses of proteins with Coomassie Blue staining.

The molecular mass for GSU0105 was measured with LC/ESI-MS. The liquid chromatography used a 2-90% MeCN linear gradient in water with 0.1% formic acid on a C18 BioBasic analytical HPLC column (Thermo Scientific) installed on an Agilent 1290 HPLC system. The eluant was analyzed with a photodiode array detector spectrophotometer and an Agilent ESI-MS 6224 TOF.

## 3. Results and discussion

### 3.1. Sequence relationship with other triheme G. sulfurreducens cytochromes

Out of 111 c-type cytochromes encoded in the *Geobacter sulfurreducens* genome, only a small subset has been characterized as the vast majority of the studies done to date have focused on the cyt c_7_ family of triheme c-type cytochromes[14–16, 28, 31, 33, 34, 45]. Similarly to the genes for the cyt c_7_ family, the gene for GSU0105 encodes three c-type heme attachment motifs (CXXCH). Gene sequence translation predicts that GSU0105 contains 93 amino acids before its processing via removal of its periplasm signaling peptide. The expected unprocessed size of the protein is similar to the sizes of unprocessed members of the cyt c_7_ family: 90 amino acids for PpcE, 91 for PpcA and PpcB, 92 for PpcD, and 95 for PpcE. Interestingly, unlike the cyt c_7_ family with six histidine residues, all involved in bis-His axial ligation of the c-type heme groups, GSU0105 has only four histidines. This suggests that at least two hemes must have mixed axial ligands. With three methionine residues present in the sequence, two of them appear to be the most likely candidates to serve as the heme axial ligands at neutral pH.

Sequence alignment of GSU0105 and cyt c_7_ family members (Fig.1) shows, that despite its similar size and the expected three c-type hemes, the GSU0105 sequence is significantly different and does not belong to the cyt c_7_ family. The residues corresponding to the N-terminal 25 amino acids of the mature sequence from PpcA, the most studied and best understood member of the cyt c_7_ family, are missing. The lost sequence corresponds to an antiparallel β-sheet (residues 1-16) followed by an α-helix (residues 17-25) in PpcA. The lost β-sheet constitutes a substantial part of the surface area in the cyt c_7_ family and forms van der Waals contacts with all three hemes. The missing α-helix contains two axial histidine ligands for Hemes-I and –III in PpcA. This sequence change implies the possibility of a substantial difference in the tertiary protein structure of GSU0105. Another place of significant sequence difference occurs at the end of an extended helical domain (residues 43-58) in PpcA, which contains the Heme-III covalent attachment site (Cys-51, Cys-53, His-54). The sequence of GSU0105 has a four residue insert near the C-terminus of the sequence suggesting the presence of an extra helical turn in this part of GSU0105. The third area of major sequence difference between GSU0105 and the cyt c_7_ family occurs immediately before the Heme-IV attachment site (Cys-65, Cys-68, His-69) in PpcA. In this position GSU0105 has an insert of 14 residues. 7 of the residues in the insert are charged, which suggests that this structural element is located on the surface of the protein. Finally, the C-terminus of GSU0105 has a six residue extension. In the cyt c_7_ family only PpcC has a notable extension of four residues. Overall, the sequence analysis suggests significant structural differences between GSU0105 and the cyt c_7_ family in *G. sulfurreducens*.

**Figure 1.**
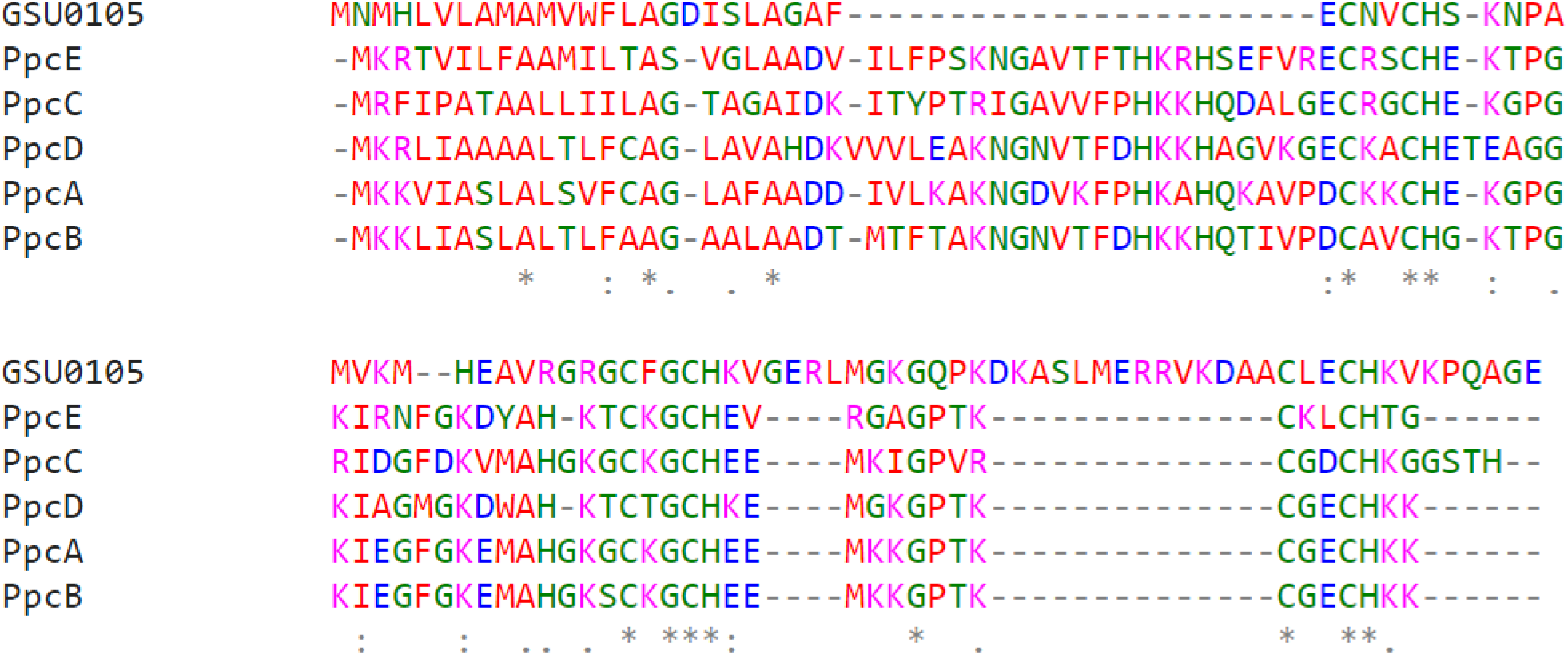
Sequence alignment of GSU0105 and all five members of the cyt c_7_ family in *G. sulfurreducens*

### 3.2. Protein expression and isolation

Previously for an unrelated project in our lab, we inserted a sequence encoding a C-terminal Factor Xa-cleavable polyhistidine tag into the pVA203 vector[31], which is widely used for expression of PpcA[20–22, 41, 46–50]. While we obtained PpcA expression levels comparable to the untagged form with the vector pVA203, and much higher yields than obtained by Londer and co-workers for their N-terminal His-tag construct [14], we have not extensively used the developed pVA203XaHT vector. This is primarily due to problems with co-purification of apo-proteins when performing metal affinity chromatography, which necessitates an additional chromatographic purification step. However, we reasoned that the presence of an affinity tag could be beneficial for expression and characterization of novel multiheme cytochromes and studies of protein-protein interactions. Therefore, we used the pVA203XaHT template to replace the PpcA gene with a gene encoding GSU0105 using restriction-free cloning[39]. We retained the N-terminal OmpA signaling sequence and the C-terminal Factor Xa-cleavable His-tag sequence. Replacement of the native *G. sulfurreducens* signaling sequence with the OmpA sequence was previously shown to significantly increase yields of c-type cytochromes expressed in *E.coli*[31]. The amino acid sequence of the construct used for GSU0105 expression is shown on Fig. 2.

**Figure 2.**
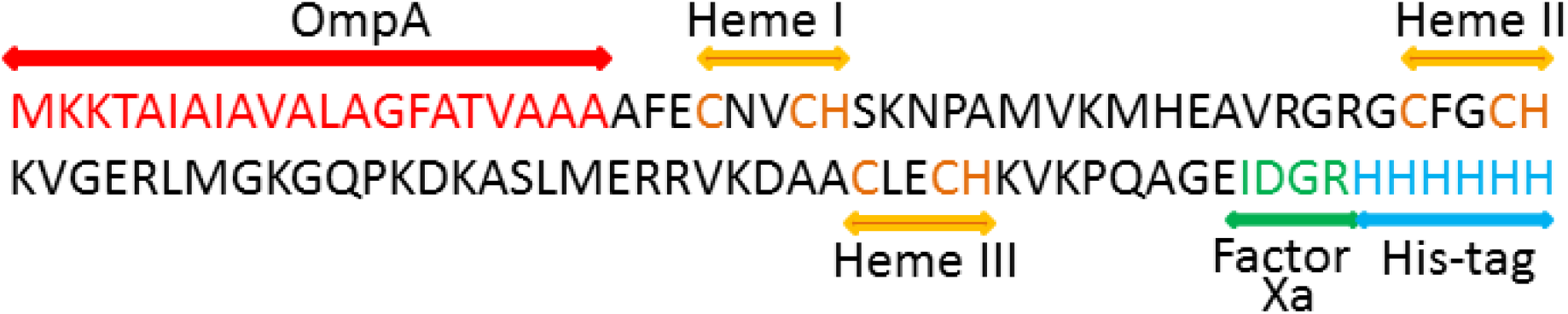
Target GSU0105 gene construct for expression in *E. coli.*

GSU0105 was isolated in two chromatographic steps. First, periplasm was applied to an equilibrated Macro S cation exchange column. A significant fraction of contaminant proteins passed through the column without binding. GSU0105 formed a clearly visible band on the column which was eluted with Buffer A supplemented with 200 mM NaCl. While the eluted protein primarily contained GSU0105, some impurities were still visible on SDS-PAGE (Fig. 3, “E1” fraction). To further purify GSU0105, we employed metal affinity chromatography as our second chromatographic step. The purified protein showed single band purity on SDS-PAGE (Fig. 3, “E2” fraction), while the flow-through and washing fractions contained removed impurities without a significant loss of GSU0105. We observed that at concentrations significantly above 100 μM, GSU0105 was prone to aggregations, especially when residual imidazole was removed. To minimize GSU0105 aggregation, we were routinely adding 20 mM cholate to stocks and protein fractions with concentrations above 50 μM or those which required long-term stability.

**Figure 3.**
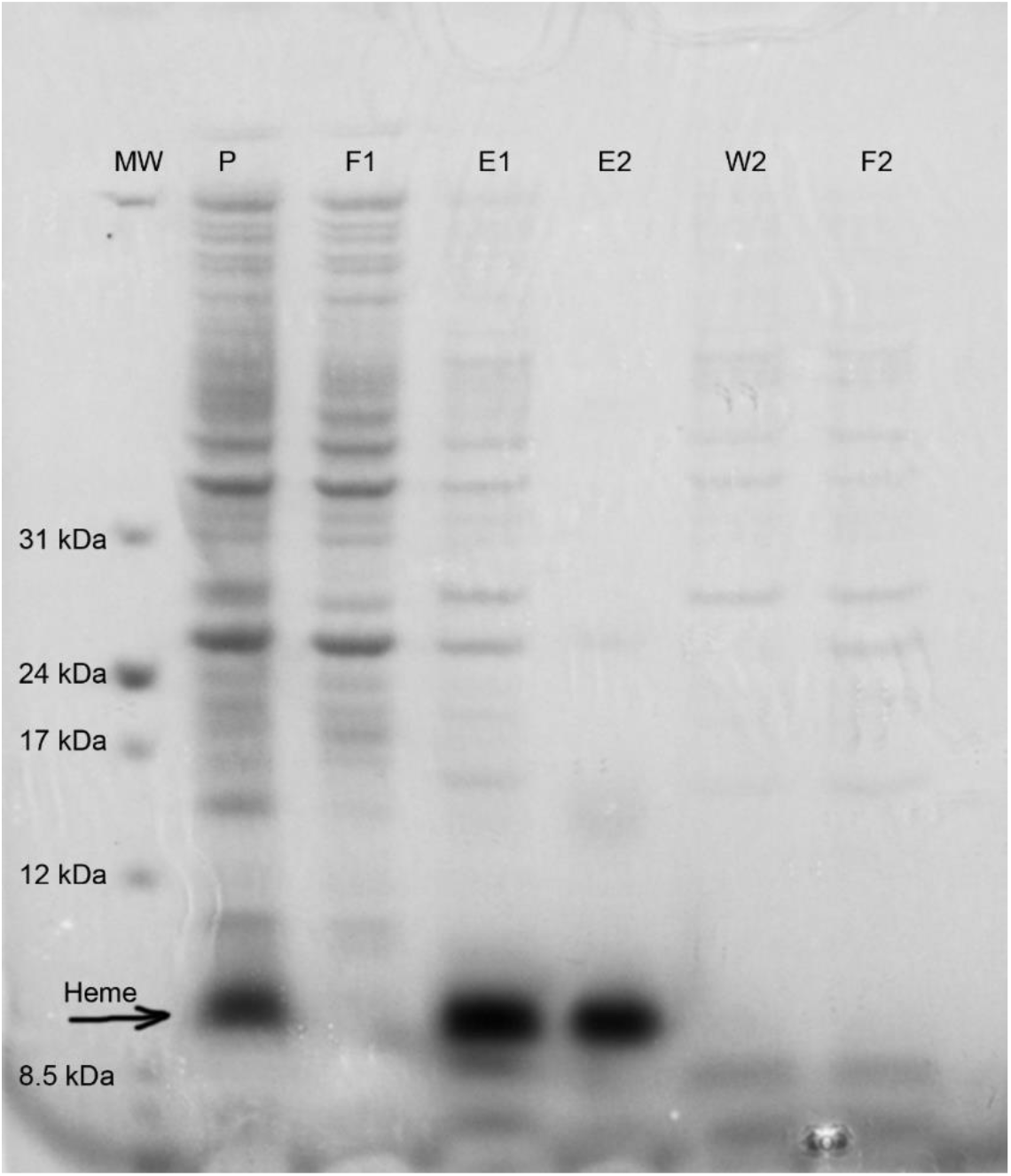
SDS-PAGE of key fractions during GSU0105 isolation. **MW**: molecular weight standards with their masses annotated in kDa. **P**: periplasmic fraction. **F1**: flow-through during binding of the periplasmic fraction to a Macro S cation exchange column. **E1**: fraction eluted with 200 mM NaCl at pH 7.5. **E2**: fraction eluted from a Ni-NTA column with 500 mM imidazole at pH 7.5. **W2**: fractions collected during washing of the Ni-NTA column. **F2**: flow-through fraction collected during binding of E1 to the Ni-NTA column. **“Heme”** denotes bands visible on the gel prior to Coomassie Blue staining.

Before Coomassie Blue gel staining we observed intense red-brown bands on non-reducing SDS-PAGE gels in the fractions corresponding to periplasm (P) and both eluants (E1 and E2). The band was located approximately half-way between the pre-stained SDS-PAGE molecular weight markers corresponding to 8.5 and 12 kDa. We attribute this band to heme-containing GSU0105, since the mass based on its amino acid sequence with the affinity tag and covalent attachment of three hemes was expected to be 10.8 kDa.

To confirm the covalent attachment of three c-type hemes and the absence of posttranslational modifications, we performed LC-ESI MS analysis (Fig. S1). We observed a typical spectrum for the protein ESI mass spectrometry with the formation of multiple ionic species with apparent masses different by 1 Da from each other. The peak with the highest intensity corresponded to +14 charge and mass of 10,849.91 Da. The expected protein mass based on complete cleavage of the OmpA signaling sequence and covalent heme attachment of 3 hemes was expected to be 10,838.88 Da for a form with a charge of +3. An acidic form of GSU0105 with 11 extra protons bound due to the presence of 0.1% formic acid in HPLC solvents should have mass of 11,849.96 Da and +14 charge. The calculated and observed masses are in an excellent agreement with each other.

Ni^2+^ leaching from Ni-NTA resin could be a concern for some biophysical characterization techniques. In our lab we developed a spectrophotometric technique to quantify Ni^2+^ leaching[51] and demonstrated that Ni^2+^ concentration during IMAC purification of GSU0105 was under 200 μM[52]. This metal contamination was removed during the buffer exchange steps targeting removal of the residual imidazole.

We routinely achieved yields above 3 mg/L of culture in BL21 (DE3) *E. coli* cells. This is significantly higher than 0.0125 mg/L in the same *E. coli* strain and 0.165 mg/L in JM109 after extensive optimization reported by T.M. Hernandez in his M.S. thesis[37]. One possible explanation for the observed difference in the yields is that the extra ten amino acids comprising the Factor Xa recognition sequence and the polyhistidine tag improved cytochrome maturation and solubility. The second factor likely contributing to the higher yield in our lab is that we were supplementing bacterial cultures with 1 mM 5-aminolevulinic acid during the induction step. However, we neither investigated this observation further nor performed significant optimization of the growth conditions as the yields obtained in our lab were comparable to the yields of the cyt c_7_ family expressed in *E. coli* and sufficient for the characterization techniques described in this report.

### 3.3. GSU0105 UV-Vis absorbance spectra

UV-Vis spectra for oxidized and reduced GSU0105 are typical for c-type cytochromes (Fig. 4). The Soret band peak position for oxidized GSU0105 is at 408 nm. Another broad absorbance peak has its maximum at 529 nm. The dithionite-reduced form of GSU0105 shows the Soret band shift to 418 nm with splitting in the range of 500-600 nm with peaks at 523 and 552 nm as expected for reduced c-type cytochromes. Interestingly, the oxidized form of GSU0105 shows a very weak charge transfer band for sulfur-iron interaction near 695 nm (Fig. 4, inset) implying that at least one of the hemes has a methionine residue as its axial ligand. This is consistent with the result reported by Fernandes[37] based on his NMR work, however, not observed in his UV-Vis spectra. Based on our data obtained with 20 μM GSU0105 (Fig. 4, inset) we can estimate the extinction coefficient at ɛ_695_ ~70 M^−1^*cm^−1^, which is significantly lower than the extinction coefficients for horse heart cyt c (1,800 M-1*cm-1) [53]. However, more than 10-fold variations of the 695 nm extinction coefficients caused by single amino acid mutations near the heme binding cavity of cyt c_2_ from *Rhodobacter capsulatus* have been previously demonstrated [54, 55] and likely indicate a more exposed to solvent heme than in cyt c.

**Figure 4.**
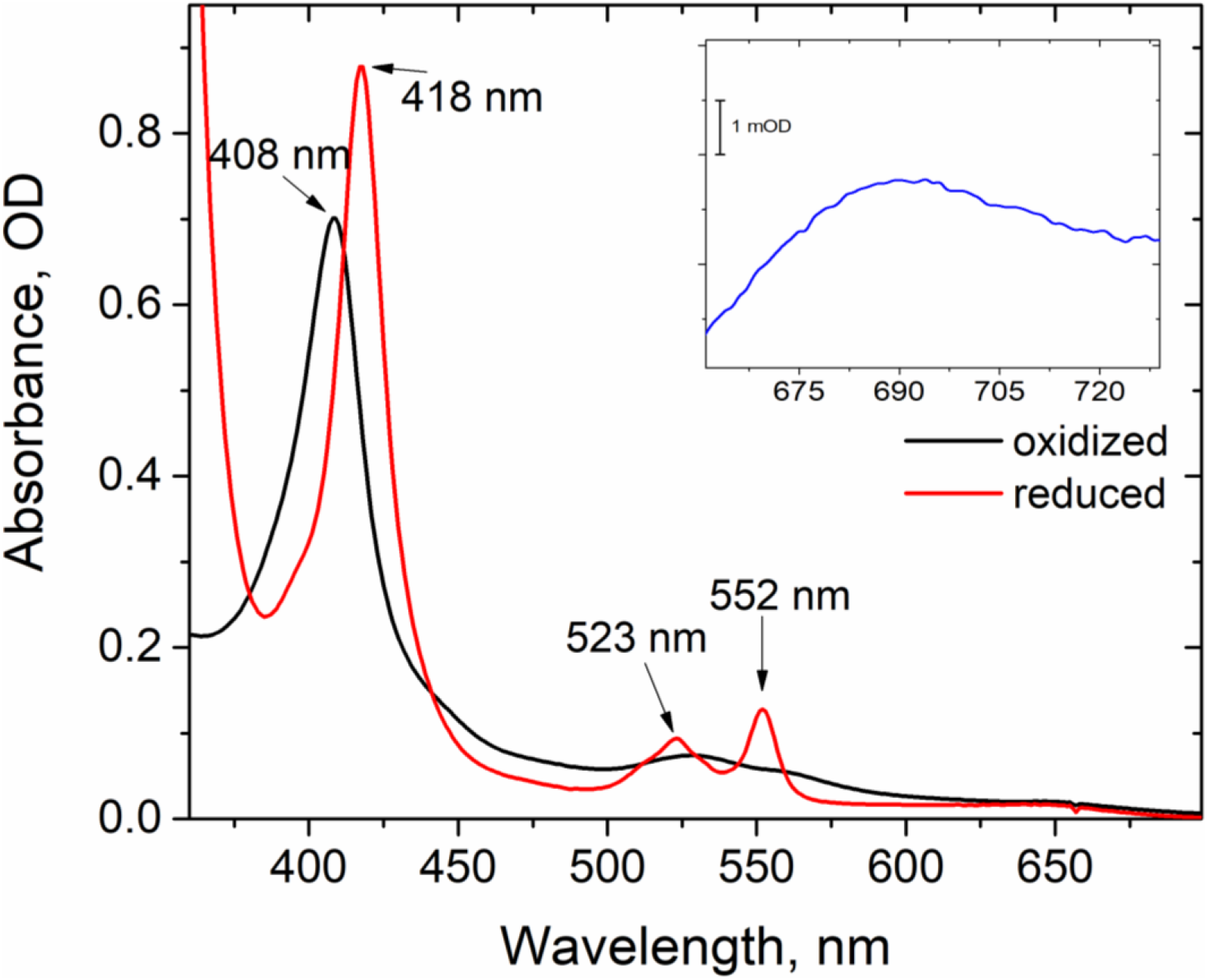
UV-Vis absorbance spectra for oxidized (black line) and dithionite-reduced (red line) 1.75 μM GSU0105 at pH 7.5. **Inset:** oxidized-*minus*-reduced spectrum of 20 μM GSU0105 near 695nm reveals a weak iron-sulfur charge transfer band.

Pyridine hemochromagen assays of heme content performed according to the procedure of Barr and Guo[43], along with an assumption that mature GSU0105 contains 3 hemes, allowed us to determine the Soret band extinction coefficient at 408 nm of oxidized GSU0105 at ɛ_408_ = 405 mM^−1^*cm^−1^.

### 3.4. Circular dichroism reveals a combination of α-helical and antiparallel β-strands in the secondary structure

Circular dichroism spectra (Fig. 5, left panel) exhibited signal consistent with the presence of α-helices and β-sheets in the protein structure. The melting curve at 210 nm (Fig. 5, right panel, ■ symbols) shows a two-step melting process with one transition observed near 40 °C and the other near 65°C. In contrast, the melting curve at 222 nm (Fig. 5, right panel, 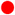 symbols) shows no significant changes until 60°C. To better understand temperature-induced changes of the GSU0105 secondary structure, we used the BeStSel algorithm to deconvolute the relative content of the secondary structure elements as a function of temperature [42]. Temperatures above 80°C were not included in our analysis due to significant protein aggregation (Fig. S2). Table S1 summarizes the relative fractions of secondary structure elements at the key transition points. BeStSel revealed that α-helical content decreased from 35.6% to 9.5% with the temperature increase from 20 °C to 60 °C (Table S1) corresponding to the first melting step observed at 210 nm. During the second melting step (60 °C to 80 °C) α-helical content further decreased from 9.5% to 4.0%. Antiparallel β-sheets showed more gradual decrease of relative content with temperature: 27.7%, 22.2%, 15.9%, and 8.9% for 20 °C, 40 °C, 60 °C, and 80 °C, respectively. In contrast, the relative content of parallel β-sheets showed increases in the range of 20-60 °C (6.1%, 16.0%, and 30.3%) and a negligible change above 60 °C. The relative fraction of turns remained stable at ~7% through the entire temperature range. Overall, the CD data revealed an unexpectedly complex structural rearrangement with temperature.

**Figure 5.**
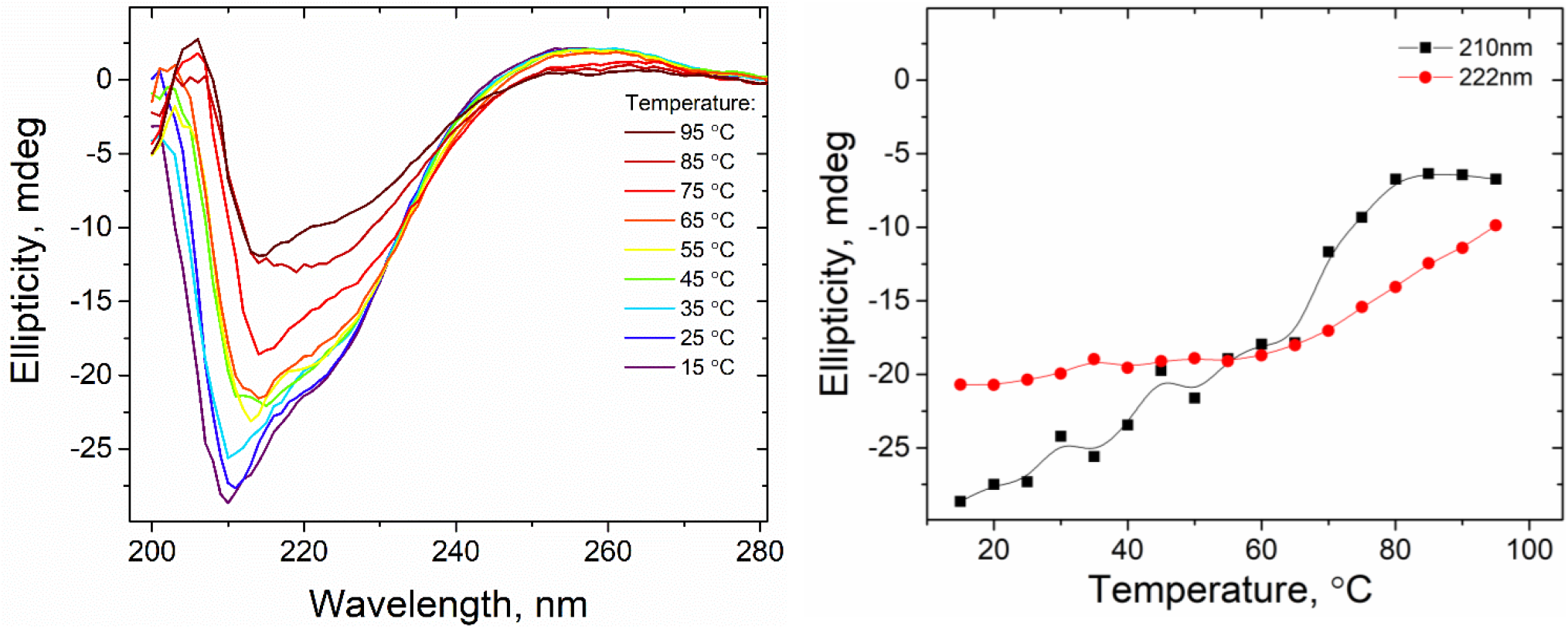
Circular dichroism studies of GSU0105. **Left panel**: CD spectra recorded with temperature increments of 10 °C. **Right panel**: CD melting curves at 210 nm (■) and at 222 nm 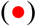

### 3.5. GSU0105 is monomeric in solution

Globular proteins are expected to produce linear SAXS Guinier plots (log I vs. q^2^) with R_g_^2^ = slope/3 [56–58]. The data for His-tagged GSU0105 (Fig. 6, left panel, 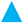 symbols) shows two linear regions from which the characteristic sizes of 14.2 Å and >68 Å can be estimated. This suggests that His-tagged GSU0105 forms elongated oligomers comprised of at least four monomers. In contrast, the Guinier plot is linear (Fig. 6, left panel, 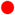 symbols) when His-tags were cleaved off by Factor Xa protease. The gyration radius for the cleaved form was estimated Rg=11.3 Å. This is consistent with globular monomeric protein with a size comparable to PpcA[41]. Displacement of native axial methionine ligands by imidazole in metal affinity chromatography preparations is a well-established phenomenon[59–61]. It is likely that in concentrated GSU0105 solutions, during dialysis or other steps for residual imidazole removal, one of the axial heme ligands in GSU0105 can be replaced by a histidine residue belonging to the His-tag of a different cytochrome molecule instead of properly refolding with the native methionine ligand in place. This process can create multimeric elongated structures with cytochromes tethered to each other through axial ligation of the heme groups by His-tags. Interestingly, with 20 mM cholate GSU0105 is present in the monomeric form even when His-tags were present. Because of the current pandemic travel restrictions, we could not test the oligomeric state of GSU0105 with SAXS in the presence of cholate. However, analytical size exclusion chromatography data show that the amounts of large aggregates (Fig. 6, right panel, ~11 min elution time) and GSU0105 dimers (~23 min elution time) are negligible in comparison with the monomeric GSU0105 form (elution time ~23 min). The calculated apparent GSU0105 mass from the size exclusion experiments based on the calibration curves shown on the inset of the right panel of Fig. 6 is 14.5 kDa. While this value is higher than 10.8 kDa expected from the sequence with three hemes bound, we note that the apparent mass is expected to be larger due to the10 amino acids from the Factor Xa recognition site sequence and His-tag protruding from the protein surface. It is also likely that there is cholate bound to GSU0105. Considering the significant cost of commercial Factor Xa, supplementing buffers with cholate during imidazole removal provides an attractive alternative.

**Figure 6.**
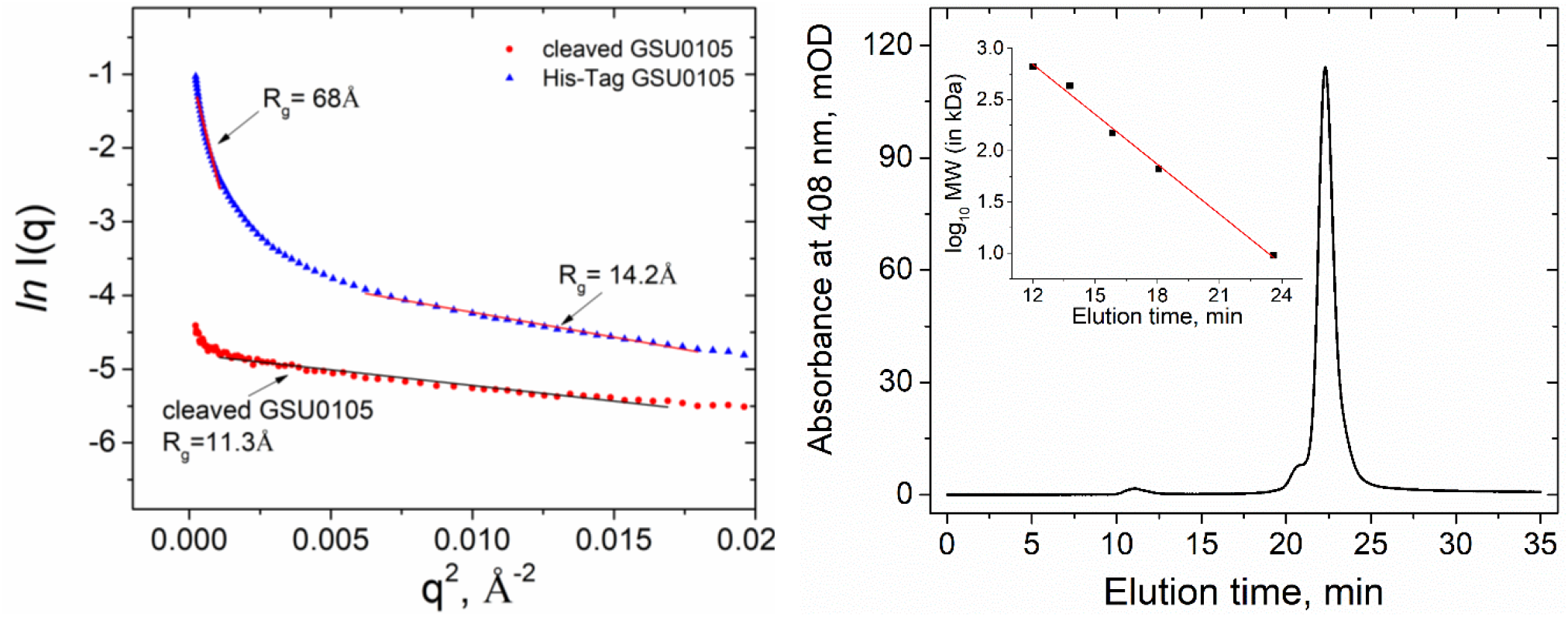
Left panel: Small angle X-ray scattering Guinier plots for His-tag GSU0105 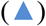 and cleaved His-tag GSU0105 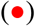 in 20 mM Tris, pH 7.5. **Right panel**: elution profile monitored at 408 nm for His-tag GSU0105 on a Superdex Increase 10/300 analytical size exclusion column in 20 mM Tris, pH 7.5, 200 mM NaCl, 20 mM cholate. **Right panel, inset**: calibration of the column with molecular weight standards.

### 3.6. GSU0105 binds to the cyt c_7_ family cytochromes

We exploited the presence of the His-tag on GSU0105 to perform affinity pull-down assays and to study GSU0105 interactions with the cyt c_7_ family members. Out of five members of the Ppc family, only PpcC has a histidine residue not buried inside of the protein and not involved in the heme axial ligation. Therefore, we did not expect any significant non-specific interaction between the Ppc family members and Ni-NTA resin. For the affinity pull-down assay, we incubated resin aliquots with ~1.3 nmol of GSU0105 (Fig. 7, red bars) or no GSU0105 (green bars) samples with 10 nmols of the respective cyt c_7_ family members or buffer. With three steps of washing with combined >100-fold dilution, we expected the residual amounts of non-specifically bound Ppc cytochromes to be less than 10% of the amount of GSU0105 bound in the “+GSU0105” samples. However, in all cases we observed higher than expected normalized residual amounts of Ppc cytochromes bound to Ni-NTA resin, which could be eluted with 500 mM imidazole: 15.9 ± 1.8% for PpcA, 32.8 ± 3.6% for PpcB, 74 ± 3% for PpcC, 72 ± 3% PpcD, 22.7 ± 4.2% PpcE. In the absence of clusters of histidine residues on the protein surfaces, this affinity to Ni^2+^ immobilized on the resin beads was unexpected and may indicate the presence of native transition metal binding sites on the surfaces of Ppc cytochromes.

**Figure 7.**
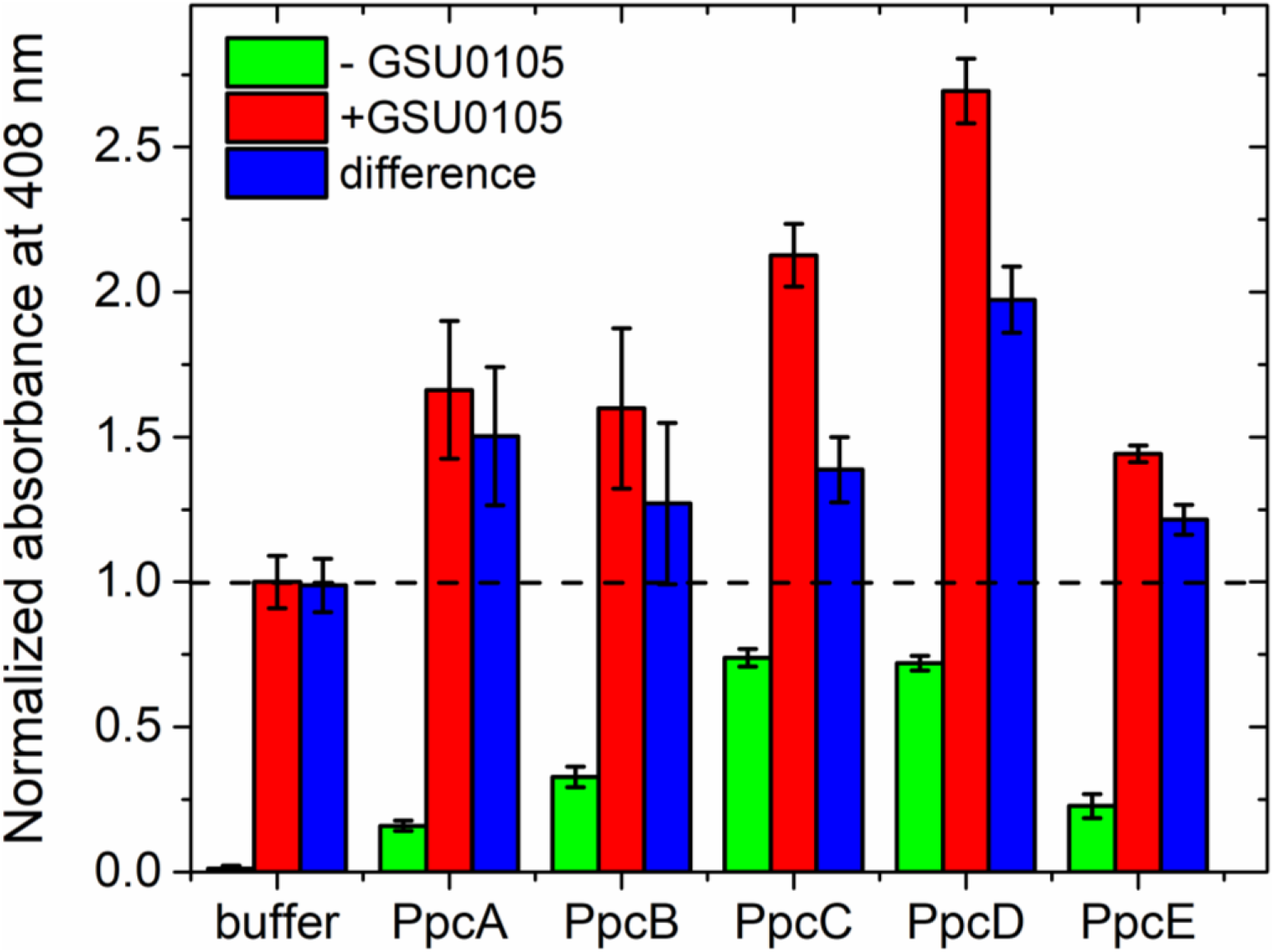
Affinity pull-down assays with GSU0105. **Red bars**: normalized heme absorbance for eluted proteins in affinity pull-down assays with GSU0105 immobilized on Ni-NTA resin. **Green bars**: retention of non-specifically bound cytochromes to Ni-NTA resin. **Blue bars:** difference between amounts of cytochromes eluted with and without GSU0105 immobilized to Ni-NTA resin. The bar heights represent the average normalized absorbance values and the error bars represent standard deviations from 3-4 independent experiments.

Despite some non-specific binding of Ppc cytochromes to Ni-NTA resin, significantly higher amounts of cytochromes were eluted for the samples with GSU0105 immobilized to the resin beads (Fig. 7, red bars). The differences in the amounts of eluted cytochromes in the presence and in the absence of GSU0105 are shown with blue bars on Fig. 7. The differences were: 1.50 ± 0.24 for PpcA, 1.27 ± 0.28 for PpcB, 1.39 ± 0.11 for PpcC, 1.97 ± 0.11 for PpcD, 1.21 ± 0.05 for PpcE, and 1.00 ± 0.09 for the buffer control experiments. Since the Soret band extinction coefficients of GSU0105 and Ppc cytochromes are similar, the normalized absorbance of 1.0 in the affinity pull-down assay would indicate the absence of protein-protein interaction and the elution of only initially immobilized GSU0105. In contrast, the normalized absorbance of 2.0 would indicate a tight 1:1 binding and elution of equal amount of GSU0105 and its binding partner. Our data demonstrates that PpcD tightly binds to GSU0105. Since GSU0105 was incubated with 10 μM PpcD and the stoichiometric amount was retained after the course of 3 washing steps spanning over 30 min, this suggests that the binding affinity of GSU0105 and PpcD is significantly tighter than K_d_ = 10 μM. Our data is less conclusive about the strength of interactions of GSU0105 with the other members of the Ppc family. However, weaker binding interactions of the other four tested cytochromes with GSU0105 are possible. We are currently working on quantifying the affinity of these interactions with isothermal calorimetry and surface plasmon resonance. We expect to present our results in a separate manuscript soon.

T.M. Fernandes in his M.S. thesis reported the results of cyclic voltammetry experiments with GSU0105 showing heme potentials: 91 ± 16 mV, −100 ± 23 mV, and −212 ± 26 mV at pH 7[37]. This range spans above and below the typical potentials of the cyt c_7_ family (e.g., −156 mV, −139 mV, and - 149 mV for PpcD)[37, 62]. Therefore, it is feasible that GSU0105 has evolved to expand the range of available redox potentials in periplasmic *G. sulfurreducens* cytochromes for more efficient linking of cytosolic and extracellular redox processes.

## 4. Conclusions

*G. sulfurreducens* is a fascinating model organism for understanding fundamental principles of bioenergetics in dissimilatory metal-reducing bacteria. Despite more than three decades of studies [8, 13, 63–65], the electron transfer pathways from cytosol to extracellular electron acceptors and their changes in response to environmental conditions are not mapped out even for Fe(III) respiration. Proteomics studies from *G. sulfurreducens* grown under various conditions have identified patterns in the changes in the proteome expression (see e.g., [35]) and draw attention to a putative triheme c-type cytochrome GSU0105 as an important link in the Fe(III) reduction pathway. In this report we have demonstrated a successful cloning and recombinant expression of GSU0105 in *E. coli* with yields over 3 mg/L of culture. This is comparable to the expression levels of the better studied members of the Ppc family [14, 16] and should enable NMR and X-ray crystallographic structural studies. ESI mass spectrometry confirmed attachment of the expected three hemes.

Despite the size and heme content similarities with the Ppc family of cytochromes, sequence analysis reveals that GSU0105 does not belong to the cyt c_7_ family. Through a combination of SAXS and analytical size exclusion experiments we found that the protein is present in the monodisperse form in solution. We note that GSU0105 appears to be prone to aggregation, which can be minimized by additions of cholate. It is currently not clear if cholate stabilization is an intrinsic phenomenon or an artifact due to the presence of the affinity tag. It is feasible that GSU0105 may interact with humic substances present in the periplasmic space and cholate molecules may have affinity for the same binding site.

While UV-Vis spectral features in the Soret and α- and β-bands are typical for c-type cytochromes, we also observed a weak charge transfer band at 695 nm suggesting that at least one of the heme groups has a methionine residue as its axial ligand. Again, this differentiates GSU0105 from PpcA-PpcE cytochromes. CD spectroscopy revealed a mixture of the secondary structure elements with a surprising conversion of α-helices and antiparallel β-sheets to parallel β-sheets at elevated temperatures. Additional work is needed to further study this phenomenon and to understand if it has any physiological significance. It is feasible that this temperature-related conformational change can lead to changes in heme redox potentials and affect GSU0105 binding to other proteins and substrates. In turn, this can result in selection of different electron transfer pathways at different temperatures.

Finally, exploiting the engineered polyhistidine tag, we observed tight binding of GSU0105 to PpcD and possibly the other members of the cyt c_7_ family in the affinity pull-down assays. These interactions need to be explored further with more quantitative techniques such as isothermal calorimetry and surface plasmon resonance. However, if verified, it would suggest that GSU0105, with its wider range of heme potentials[37], evolved to shuttle electrons in the periplasm from the Ppc family to other substrates, including the outer membrane cytochromes involved in Fe(III) reduction.

## Acknowledgments

The authors are grateful to Prof. D.R. Lovley (University of Massachusetts-Amherst) for his generous gift of genomic *G. sulfurreducens* DNA. This research was funded by the National Science Foundation, grant numbers RUI MCB-1817448 and REU CHE-1757874. X-ray scattering experiments were carried out at beamline 12-ID-B of the Advanced Photon Source, an Office of Science User facility operated for the U.S. Department of Energy (DOE) Office of Science supported at Argonne National Laboratory by the U.S. DOE under Contract DE-AC02-06CH11357. The authors gratefully acknowledge help of Dr. Xiaobing Zuo and staff of Sector 12 of the Advanced Photon Source. Finally, the authors acknowledge help and support from Dr. P.R. Pokkuluri (Argonne National Lab and Auburn University).

