## Supplemental figures and tables for "Isolation and biophysical characterization of GSU0105, a triheme c-type cytochrome from *Geobacter sulfurreducens*"

### Supporting information

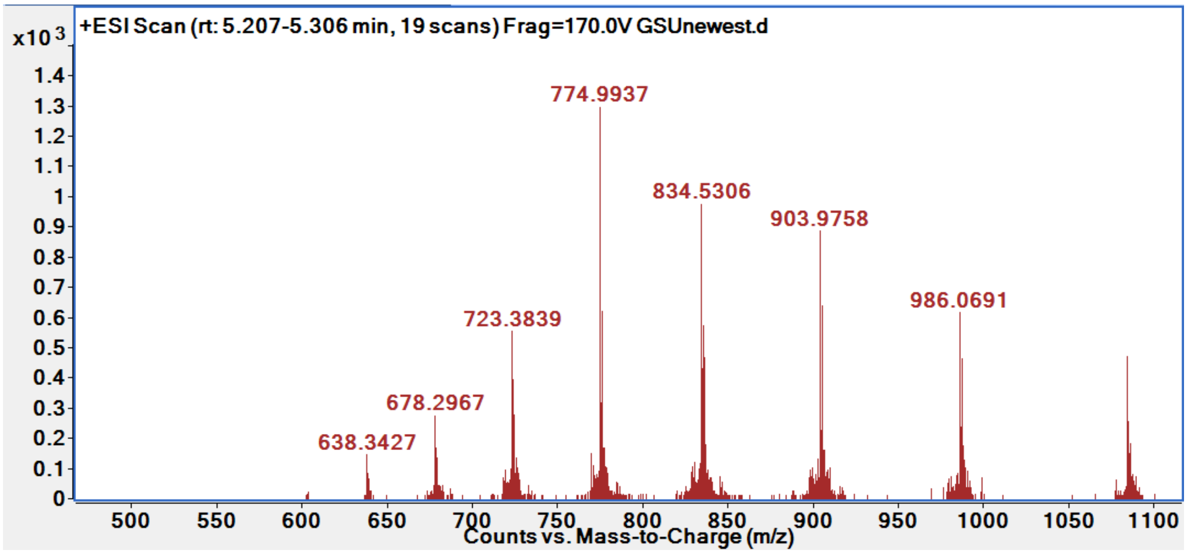


**Figure S1.** ESI-MS spectrum of GSU0105 in 0.1% (v/v) formic acid.

| | 20°C | 40°C | 60°C | 80°C |
|---|---|---|---|---|
| α-helix | 35.6% | 18.8% | 9.5% | 4.0% |
| Antiparallel β -sheet | 27.7% | 22.2% | 15.9% | 8.9% |
| Parallel β -sheet | 6.1% | 16.0% | 30.3% | 27.4% |
| Turn | 7.4% | 7.4% | 5.6% | 9.0% |
| Others | 23.2% | 35.6% | 38.6% | 50.7% |

**Table S1.** Secondary structure of GSU0105 as a function of temperature.

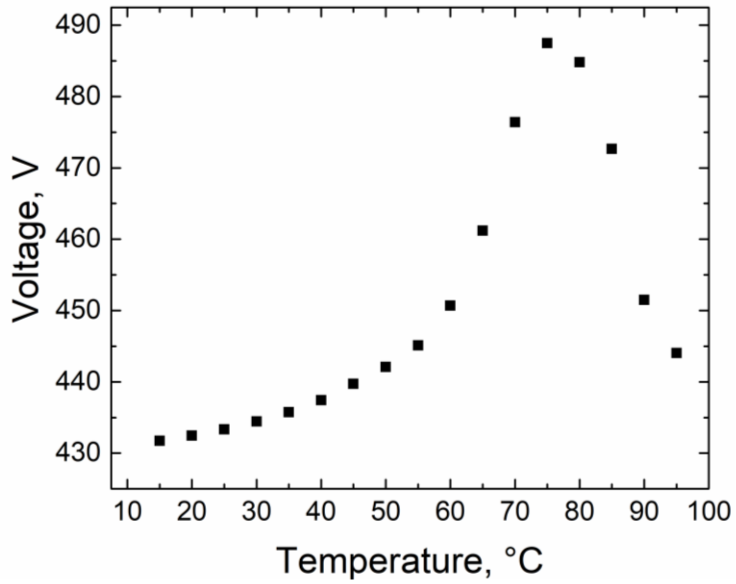


**Figure S2.** Turbidity of GSU0105 solution at 222 nm as a function of temperature.
